## Supplementary Information for "Factors influencing taxonomic unevenness in scientific research: A mixed-methods case study of non-human primate genomic sequence data generation"

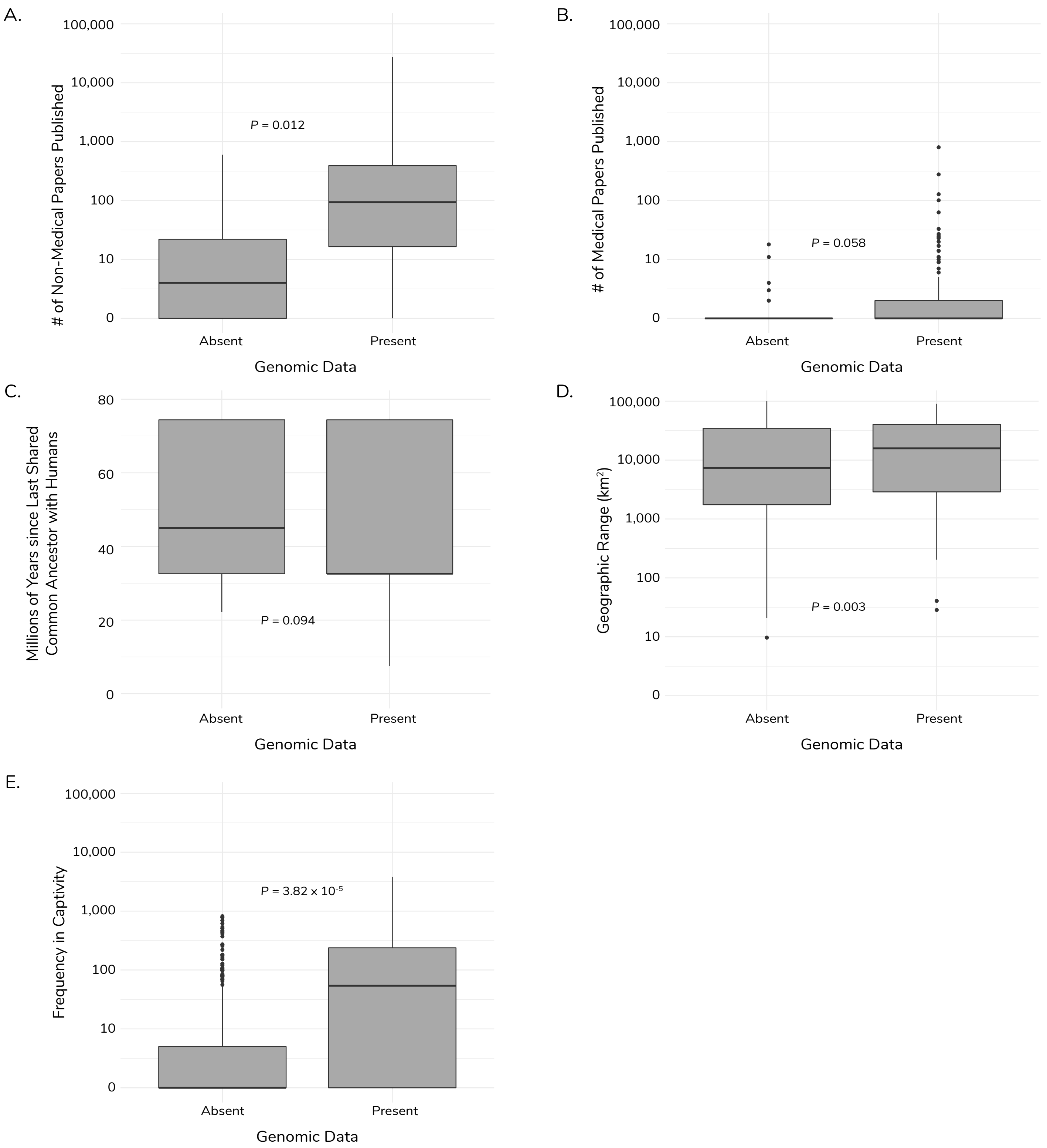


**Supplementary Figure 1. Boxplots comparing species with and without any genomic data for each tested variable.** The box extends from the first to the third quantile. The horizontal line within each boxplot represents the median value. The whiskers represent the lower and upper extreme value limits. The black dots represent outliers.

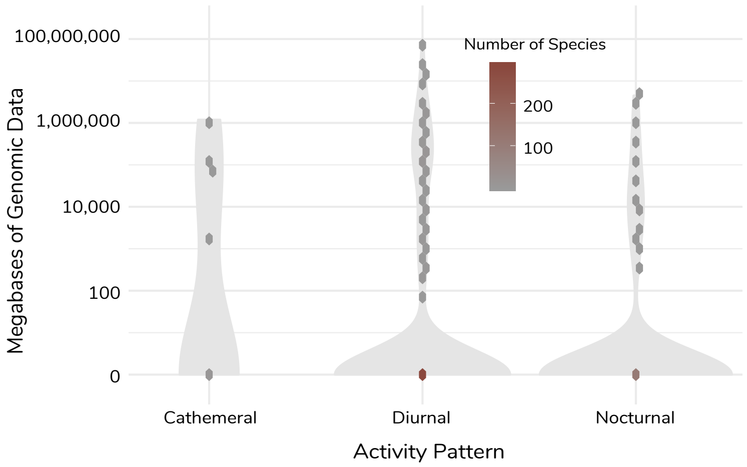


**Supplementary Figure 2. Violin plots of per-species genomic data by activity pattern.** Violin plot width corresponds to the density of species, which is also depicted via heatmap.

| Genus Grouping | Collapsed Genera |
|---|---|
| Callicebus | Callicebus, Plecturocebus |
| Callithrix | Callithrix, Cebuella, Mico |
| Cebus | Cebus, Sapajus |
| Cercopithecus | Cercopithecus, Chlorocebus, Erythrocebus, Allochrocebus |
| Galago | Galago, Galagoides, Otolemur, Schuirocheirus |
| Lagothrix | Lagothrix, Oreonax |
| Papio | Papio, Rungwecebus |
| Saguinus | Saguinus, Leontocebus |
| Trachypithecus | Trachypithecus, Semnopithecus |

**Supplementary Table 1. List of paraphyletic generic names found in genomic databases that were collapsed into a single genus for construction of Figure 1.** The single genus names listed in the first column were used in Figure 1 (denoted by asterisks in the Figure) to represent data combined for multiple genera now believed to be paraphyletic, listed in the second column.

| List of Interview Questions |
| --- |
| Q1. Why did you decide to do work on xx species? Was your lab already working on this species, or was this the first time you did a particular project on this species? |
| Q2. What motivated you to do this study/start working with this species? |
| Q3. Were there factors that made studying your select-ed primate species easier than it would have been to study other potential species? If so, what were they? |
| Q4. Were there factors that made studying your select-ed primate species more difficult than it would have been to study other potential species? If so, what were they? |
| Q5. Did you face any additional challenges in working with your selected species? |
| Q6. Did you want to say anything additional that you didn't get the opportunity to say already? |

**Supplementary Table 2. List of interview questions.** Interview questions were asked in the order presented in this table. At times, additional probing questions were asked regarding the answers provided for each question. However, overall, each interview followed the structure presented here.

| Deposit Name | Total Data (Mb) |
|---|---|
| Cebus sp. | 402 |
| Gorilla | 210,516 |
| Macaca | 422,040 |
| Microcebus | 586 |
| Papio anubis x Papio hamadryas | 35,584 |
| Papio anubis x Papio cynocephalus | 898 |
| Papio anubis x Papio ursinus | 1,162 |
| Papio kindae x Papio cynocephalus | 1,932 |
| Papio kindae x Papio ursinus | 17,119 |
| Primates | 1,176 |
| Rhinopithecus | 480,606 |
| Unidentified | 242 |
| Unidentified monkey | 9,546 |
| **Total** | 1,181,809 |

**Supplementary Table 3. SRA deposits omitted from genomic dataset.** A complete list of the SRA deposits that were removed from the dataset (see Methods), including the total amount of genomic data under each deposit name. Deposits that were not identified at the species level or that came from known species-hybrid individuals were omitted.

| Data | Test | Distribution | Variables[1] | AIC | p-value | $r^2$ |
|---|---|---|---|---|---|---|
| Whole dataset | Logistic regression | Binomial | Non-medical publications + Relatedness to humans + Medical papers published + Geographical range + Frequency in zoos + IUCN Red List Status + Activity pattern | 346.3 | N/A | N/A |
| Whole dataset | GLM | Gaussian | Non-medical publications + Relatedness to humans + Medical papers published + Geographical range + Frequency in zoos + IUCN Red List Status + Activity pattern | 1659.3 | $<2.2\times10^{-16}$ | 0.40 |
| Species with genomic data | GLM | Gaussian | Non-medical publications + Relatedness to humans + Medical papers published + Geographical range + Frequency in zoos + IUCN Red List Status + Activity pattern | 293.26 | $6.16\times10^{-8}$ | 0.40 |

Note:
[1] "+" indicates the function used for the variables within each model.

**Supplementary Table 4. Analytical models performed.** A list of all models used within the study, including data used, test performed, distribution of data, variables included, AIC values, p-values, and $r^2$ values where applicable.
